# Are you represented? Subjective vs objective skin color determination for healthcare and research purposes

**DOI:** 10.64898/2026.04.13.718177

**Authors:** Kerry Setchfield, Suvvi K. Narayana Swamy, Elena J.Setchfield, Stephen P. Morgan, Michael G. Somekh, Amanda J Wright

## Abstract

Despite questionable accuracy, subjective methods to categorize skin color are heavily relied upon in research and medicine. Objective skin color determination is expensive requiring specialized instrumentation and interpretation. We compare three subjective approaches, i) Fitzpatrick Skin Type Scale (FST), ii) Pantone SkinTone Guide (PST) and, iii) Monk Skin Tone Scale (MST), with objectively measured skin color from a spectrophotometer in 87 volunteers to understand the limitations of each method. In agreement with others, we show that the popular FST questionnaire correlates poorly with the objective approach. However, PST color swatches provide good correlation with spectrophotometer measurements. PST consists of 110+ swatches that are inexpensive and easy to use, however, similar to other reports, the volunteers found the number of swatches overwhelming and/or excessive. We found that the recently introduced MST is not representative of reality with only 3 of the 10 color groups representing our volunteers and published populations of volunteers. In future, we propose using 9 color swatches to split the spectrum of human skin color into 10 groupings (Nottingham Skin Categories - NSC) that are representative of the global population. This new approach would be easy to implement and inexpensive in research, healthcare and cosmetics settings, and maps directly to objective, quantitative, measures taken with a spectrophotometer. For the testing and development of new optical devices, NSC would provide increased comparability between studies and ensure studies are representative of local/global populations. In the clinic NSC would be useful for dermatology, photodynamic therapy and dosage assessment for topical medicine, for example.

## INTRODUCTION

Optical medical technologies form an increasing and significantly large fraction (∼ 20%) of the global medical technologies market, being valued in excess of $120 billion USD per year in 2023 [1, 2]. These technologies include otoscopes and dermatoscopes used in clinics, more advanced technologies like lasers for surgery or cancer treatment, and imaging tools like Optical Coherence Tomography (OCT). They are used broadly across medical settings as well as in the home where medical wearables and smartwatches are becoming increasingly popular. In all cases, these technologies require the interaction of light with human tissues, such as skin, with the nature of these interactions being used to provide information about the tissue. Technologies often require calibration to the optical properties (including absorption and scattering coefficients) of the tissues being assessed. When it comes to imaging/treating skin conditions the majority of optical medical technologies are tested on fair skin[3]. During the research and testing stages of optical medical devices for example, only 1% of studies evaluate the effect of skin color, and where skin color has been established, most studies are carried out in light skin[4]. The negative consequences of this were made clear during the COVID 19 pandemic when pulse oximeters, which use two different wavelengths to determine blood oxygen saturation, over-estimated the oxygen saturation of patients with dark skin leading to poorer health outcomes in patients from this group[5–9]. Skin color has also been shown to lead to different outcomes for other medical wearable technologies, medical imaging and photodynamic therapies[4]. Recent publications urge increasing subject diversity in optical medical technology research and accurate reporting on any effect skin color has on results [4, 10, 11]. In April 2023, the UK NHS Race and Health Observatory released a report stating that there should be a review of all medical devices to determine suitability for use for people with dark skin and that this should be ‘sufficiently evidenced by manufacturers before devices receive market approval’[12]. More recently the FDA has updated its recommendations for manufacturers of pulse oximeters to improve clinical study design and device evaluation across the range of skin tones[13]. To address these requirements, a standardized, easy to use, and inexpensive method is needed to accurately determine and classify skin color.

Skin color is currently determined either subjectively, by comparison with a scale or questionnaire, or objectively using an optical device such as a spectrophotometer. Multiple subjective skin color assessment tools have been developed to determine an individual’s skin color, with the Fitzpatrick Skin Type Scale (FST) being the most commonly used today despite being shown to be inaccurate, particularly for dark skin[14, 15]. This scale assesses skin color via a questionnaire based on an individual’s reaction to UV light. Other less commonly used subjective scales include Genetico-Racial Skin Classification, Lancer Ethnicity Scale, Roberts Skin Type Classification System and the Von Luschan chromatic scale[16–19]. Like the FST, some of these include self-reporting of effects of light on skin, however, unlike the FST, they are generally used for determining the potential side effects of cosmetic laser treatments on skin. The Pantone SkinTone Guide (PST) consists of a color chart with 110 shades that are compared to the skin to obtain an understanding of an individual’s skin color and to match products, such as prosthetics, to the individual’s skin tone. Recently, the Monk Skin Tone Scale (MST) has been developed in conjunction with Google to allow representation of skin tones for computer vision applications[20]. The MST has also been suggested for use in healthcare as a better, more representative, determinant of skin color than FST [21]. The utility of the FST, PST and MST will be described in more detail in sections 1.1-1.3. Regarding objective skin color measurement, spectrophotometers and colorimeters are most used in research. Colorimeters measure skin color using red, green and blue filters mimicking human vision[22]. Spectrophotometers provide a more detailed, accurate and perceptually uniform understanding of skin color than a colorimeter and quantify skin color using an individual typology angle (ITA), see section 1.4[23].

In 2013, Del Bino and Bernerd assessed 3,500 women from various geographical areas and determined skin color objectively using a chromosphere and the ITA scale (*24*). The Del Bino study aimed to understand the biological consequences of ultraviolet radiation and did not compare subjective and objective methods for determining skin color. These authors suggested a 6-group objective color categorization system, based on ITA, to replace the FST. This consisted of 2 groups, ‘brown’ and ‘dark-brown’ to represent dark skin[24]. A recent paper by Lipnick et al (2025) compares three subjective methods, the Von Luschan (which comprises of 36 opaque glass tiles that volunteers can match to their skin color), a printed color representation of FST, and a professionally printed MST, with two objective measurements using a spectrophotometer and a colorimeter, in a study of 789 volunteers [25]. They proposed a new cut-off to the ITA scale to better reflect real world populations and people with dark skin. Because the thresholds for the groups described by Del Bino and Bernerd were considered too light by Lipnick et al., (2025), the authors have suggested further splitting these groups into 4 groups to represent ‘brown’, ‘intermediate dark’, ‘dark’ and ‘very dark’[25].

In this manuscript, we compare three subjective skin color scales, FST, PST, and MST, with skin color determined objectively using a spectrometer and expressed as an ITA value, in a cohort of 87 volunteers. Similar to others, we highlight the limitations and lack of correlation of the FST subjective classification with the objective ITA measurements, however, we show that the PST correlates well with objectively measured ITA. Nonetheless, volunteers commented that the large number of PST swatches was overwhelming/excessive. Collating the ITA volunteer data from Lipnick et al., and Del Bino and Bernerd with our data, we compare a highly diverse dataset representative of a wide range of skin colors with the range and limits of the PST and MST scales. To address the issue of the number of PST swatches we propose a new set of 9 skin color swatches (Nottingham Skin Categories – NSC) with characterized ITA values as an easy to implement, inexpensive, route to determine skin color, replacing current reliance on FST. We map these NSC swatches directly to ITA readings and propose using them to divide skin color into 10 evenly distributed groups in a range that better represents people with dark skin as described in Lipnick et al. This new NSC scale has the potential to standardize skin color classification, increase comparability amongst different research teams, and ensure diversity in device testing and design.

### 1.1 Subjective: The Fitzpatrick Skin Type Scale

The Fitzpatrick Skin Type Scale (FST) was created in 1978 to characterize an individual’s response to UV light for phototherapy treatment of psoriasis[14]. It determines the individual’s risk of burning or tanning and has since been commonly used to determine their skin cancer risk[26]. It relies on self-reporting in response to questions regarding genetic disposition (e.g. eye, hair and skin color, and number of freckles) and reaction to sun exposure (burning and tanning potential, and sun sensitivity) using a simple questionnaire (Figure A1). The scale was initially created for white skin, classified into 4 groups (FST I-IV), but was adapted in 1988 to include 2 further groups encompassing individuals with dark skin (FST V and VI)[27]. The ease of use of the FST scale has contributed to its widespread use in healthcare and healthcare-associated research. However, the subjectivity of the scale and the imprecise classification of the dark-skinned population has been widely acknowledged as an issue. The questions used to categorize individuals into FST groups are not applicable to people with dark skin who often do not associate tanning with their skin and burning is often unseen. Where used in healthcare for assigning phototherapeutic regimes, inaccurate assessment of dark skin can lead to epidermal injury[28]. Reports on responses to laser resurfacing of skin for treatment of acne scars, wound healing and other cosmetic purposes have all relied upon FST classification of patients, and showed wide variability in the extent of treatment induced skin trauma [29]. Goldman suggests that this response variability is due to inconsistency within FST group classifications[29]. In addition, use of the FST to categorize skin color can lead to the perception that people with dark skin will not suffer skin cancers; despite skin cancers being less frequent in dark skinned people than those with light skin, the morbidity tends to be higher as a consequence of later presentation and diagnosis due to an illusion of safety associated with being placed in FST V or VI[26]. A recent study looking at the accuracy of skin cancer detection in images from different FST groups suggests that this method for categorizing skin color can lead to AI training disparities for skin cancer diagnosis[30]. Available data sets are skewed towards light skin and disease diagnosis accuracy varies across different FSTs. This suggests that use of an objective method for determining skin color would be more useful for this form of AI training and that diverse skin tones need to be incorporated into training sets to lessen skin color bias.

### 1.2: Subjective: The Pantone SkinTone Guide

The Pantone SkinTone Guide (PST) is a visual reference guide consisting of 110 skin tone matched colors (recently updated to include a further 28 colors) printed on cards that can be compared with an individual’s skin [31]. In the main, this guide is used in the beauty and product design industry for color matching, however, it has also been used in healthcare for matching prosthetics to the surrounding skin. Because the guide is visual, to ensure consistency in perception, it provides the best skin tone matches under standardized D65 (6500K) daylight conditions. Skin color is represented by 4 characters that represent hue, red/yellow undertone and the darkness (Figure S2)[31].

Importantly, in a study of skin color of 61 healthy volunteers from southeastern US which compared PST with colorimetry values, PST darkness values were shown to correlate with melanin values, suggesting this is a valid method for determining skin color [32]. The PST guide has been proposed as a standardized clinical tool for color matching skin allotransplants/grafts[33]. It has also shown utility for monitoring the lightening efficacy of laser treatment for axillary hyperpigmentation[34].

### 1.3: Subjective: The Monk Skin Tone Scale

The Monk Skin Tone Scale (MST) was developed in 2023 by Ellis Monk in collaboration with Google to address equality and representation in machine learning tools, including facial recognition software, and has been incorporated into online image filter and searching algorithms[20, 35]. The aim of this scale is to replace the FST which has traditionally been used for machine learning to increase diversity and therefore product efficacy across a range of skin tones. The MST was originally created for use in social sciences and was designed to represent human skin color in North and South America. For optical medical device testing, the MST is used to ensure that devices are tested on the full spectrum of skin tones.

The MST has been suggested as a useful alternative to the FST in clinical and research settings[36]. It consists of 10 skin tone levels for visual comparison with an individual’s skin but has yet to be routinely used in dermatology (Figure S3). However, since a printed version of the MST is required for research and clinical studies, printing differences may lead to inconsistent or incorrect classification of skin tone and therefore potential research and clinical risks[37]. For computer vision and machine learning purposes, light source affects accuracy and classification of skin tones in the lightest and darkest skin tones (MST levels 1-3 and 8-10 respectively). MST skin tone levels 4-7 can be difficult to classify, independent of light source (i.e. warm/cool white fluorescent/LED/halogen sources), implying that more work is required to enhance training data[38].

### 1.4: Objective: Spectrophotometric determination of skin color

Spectrophotometers are non-invasive, objective and user-independent tools that provide spectral reflectance information regarding skin color. Illumination is via a white light source (e.g. a tungsten or xenon lamp for visible light) and using a diffraction grating, detected light is split into individual wavelengths across a spectral range (typically 400 – 700 nm)[39]. A color matching function (typically, 10-degree standard observer) allows the raw spectral data to be converted to tristimulus (XYZ) colors which represent a device independent perceived color. This can then be converted to a perceptually uniform color space, known as the CIELAB uniform color space using equations described by Ly et al [22]. Because spectrophotometers measure skin color objectively and accurately, they are helpful in describing it in an unbiased manner. They have been used in clinical applications including assessing pigment changes associated with disease, chemotherapy and surgery as well as for evaluating donor skin in transplants[33, 40–42].

The CIELAB color space was developed in 1976 by the International Commission on Illumination (CIE)[43]. Color is expressed as L* for luminance, a* as the amount of red/green and b* as the amount of blue/yellow each given as a numerical value and changes which correspond to perceived color change. These changes are standardized across color capture devices so that measured color is device independent. L* values range between 0 (black) and 100 (white) and positive a*/b* values being red/yellow colors respectively and negative a*/b* values being green/blue colors respectively (Figure S4a). A standard D65 illuminance utilised and represents average daylight including UVA and longer wavelength UVB. From the L* and b* values the Individual Typology Angle (ITA) can be calculated, and this has been recommended for further classifying skin into 6 objective skin color groupings as described by Del Bino and Bernerd[24, 44]. ITA values are calculated from L* and b* values only (a* values are not used) because these are best correlated with the visual assessment of the skin color and shown to relate to melanin content of the skin[45].

Originally ITA groups consisted of 4 skin color bands created to correspond with the 4 visual color groupings that were usually used to select Caucasian individuals for assessment of skin protection products[44]. The groups were categorized as ‘very light’ with an ITA > 55°, ‘light with an ITA > 41°, ‘intermediate’ with ITA > 28° and ‘tan’ with ITA > 10° (Figure S4b and c). Two further skin color groups were later suggested by Del Bino *et* al; in 2013 to represent dark skin and defined as ‘brown’ with an ITA > -30° and ‘dark brown’ where ITA is < -30°[46]. The ITA is calculated as follows: ITA° = (ArcTan((L* −50)/b*)) × 180/π[44].

The melanin quantity in the skin correlates with L* and the ITA value, with larger values of each relating to lighter skinned individuals. A 2013 study of 3500 individuals showed the skin color spread with geographical area and the wide diversity of skin color within these regions[24]. Caucasian, Russian, Chinese and Japanese skin tended to fit within the light - tan skin categories, although Russian volunteers tended to be more skewed towards the light category. Hispanic and Thai skin tended to fit into the intermediate and tan skin color groups, while Indian skin was mainly intermediate – brown. The darkest skin was found amongst the African cohort which was mainly categorized as being tan – dark brown. Interestingly only one individual from the measured cohort was found to have skin fitting into the very light category.

### 1.5: The pros and cons of subjective and objective methods for determining skin color

By way of summary, the advantages and disadvantages of using the FST, PST, MST and spectrophotometers for determining skin color are shown in table 1.

**Table 1.** Advantages and disadvantages associated with skin color determination using FST, PST, MST and spectrophotometry.

|  | Skin colour assessment tool | Pros | Cons |
| --- | --- | --- | --- |
| Subjective methods | Fitzpatrick Skin Type Scale | Ease of use<br>Useful in light skin for understanding skin cancer risk | Subjective<br>Limited accuracy in people with dark skin<br>Contributes to healthcare disparities<br>Inaccurate<br>Ill-defined descriptors in questionnaire<br>Commonly used in research where skin color is important |
|  | Pantone SkinTone Guide | Does not require an understanding of interaction of light with skin<br>Representative and comprehensive<br>No software, calibration or extensive user training required<br>Correlates well with melanin content<br>Relative ease of use<br>Low cost (~150USD) | Requires correct lighting for reliable color matching<br>Observer dependent<br>Relatively subjective |
|  | Monk Skin Tone Scale | Simple to use: consistent color perception without requirement for specific instrumentation<br>Representative<br>Open source<br>Useful in computer vision applications | Printed variability due to paper color and quality and printer calibration leading to inconsistent skin tone classification<br>Limited skin tone range<br>Subjective<br>Not created for healthcare applications |
| Objective method | Spectrophotometer | Objective, accurate and user independent<br>Spectral data can be converted into perceptually uniform color space<br>Enables unbiased healthcare and research | Expensive (~10,000USD)<br>Requires specialised hardware and software |

## RESULTS

### 2.1: Volunteer Cohort

A total of 87 volunteers were recruited for skin color analysis and method comparison. 56% of our volunteer cohort were female at birth and 44% male. The mean age was 31 years (range of 18-71 years and median of 29 years) and the mean temperature of the laboratory was 20.6×C (range of 16.6 ×C -24.3 ×C).When categorized objectively as ITA groups 12.6%, 35.6%, 25.3%, 14.9% and 11.5% of volunteers fell into the ‘light’, ‘intermediate’, ‘tan’, ‘brown’ and ‘dark brown’ categories respectively. We were unable to recruit any volunteers in the ‘very light’ ITA skin color group (i.e. ITA > 55°). This is unsurprising given that amongst a volunteer cohort of 3500 studied by Del Bino and Bernerd only 1 person (<0.03%) was designated ‘very light’[45].

Using the ITA skin color groups described by Del Bino and Bernerd and in section 1.4, ITA ‘very light’ has a bin size of 35°, ITA ‘light’ = 14°, ITA ‘intermediate’ = 13°, ITA ‘tan’ =18°, ITA ‘brown’ = 40° and ITA ‘dark brown’ = 60°[24]. The ITA bin sizes for ‘very light’ and ‘dark brown’ extent to the theoretical limits for the ITA formula. The ITA range of volunteers in FST III, IV and V is similar (with FST V having the greatest ITA range) and includes volunteers who have objectively measured skin colors ranging from ‘light’ to ‘dark-brown’ and highlighting the subjectivity of the FST scale (table 2). The ITA ranges of volunteers in FST I and II were objectively measured as being in the ‘light’ to the lighter third of the ‘tan’ ITA groups. This confirms the original creation of the FST scale for people with light skin. Most volunteers whose skin was objectively measured to be in the ‘light’ ITA category had FST III skin, whereas most in the ‘intermediate’, ‘brown’ and ‘dark brown’ ITA categories had FST IV skin highlighting the subjectivity of the FST scale.

**Table 2.** Volunteer ITA range within each (A) ITA and (B)FST skin color group. The predominant questionnaire-resultant FST group for each ITA group is shown. Note that none of the volunteer cohort was designated as having ITA ‘very light’; one volunteer was determined to be FST VI using the FST questionnaire.

| (A) |  |  |  | (B) |  |  |
| --- | --- | --- | --- | --- | --- | --- |
| ITA group | Number of volunteers | Measured ITA° range | predominant FST color group | FST | Number of volunteers | Measured ITA° range |
| Very Light | 0 | - | - | I | 4 | 45.25 - 23.34 |
| Light | 11 | 52.98 - 41.89 | III | II | 8 | 49.10 -28.66 |
| Intermediate | 31 | 39.50 - 28.46 | IV | III | 21 | 52.98 - -52.69 |
| Tan | 22 | 27.67 - 10.03 | IV | IV | 37 | 38.23 - -65.67 |
| Brown | 13 | 9.93 - -28.96 | V | V | 15 | 42.08 - -65.65 |
| Dark Brown | 10 | -33.14 - -65.67 | IV | VI | 1 | -52.49 |

### 2.2: Spectral spread of volunteer skin color

Spectral and L*a*b* data was collected for each volunteer using the Konica Minolta CM700d spectrophotometer between 400 - 700 nm. In addition, the spectrum and L*a*b* coordinates for each PST swatch chosen to represent an individual volunteer’s skin color was measured using the spectrophotmeter. As previously described, only the L* and b* values are required for calculating an individual’s skin color, because these values provide best correlation with melanin content. Therefore, using the L* and b* coordinates, the ITA for both objectively measured and chosen PST for each individual were calculated. When color coded for measured PST associated ITA volunteers in the ‘tan’ ITA bin assess their color using the PST swatches as being a range of colors from ‘light’ to ‘brown’ (Figure 1b). The subjectivity of the FST is exemplified in Figure 1c; volunteers are categorized into FST groups that do not correlate with the objectively measured ITA color groups.

**Fig. 1.**
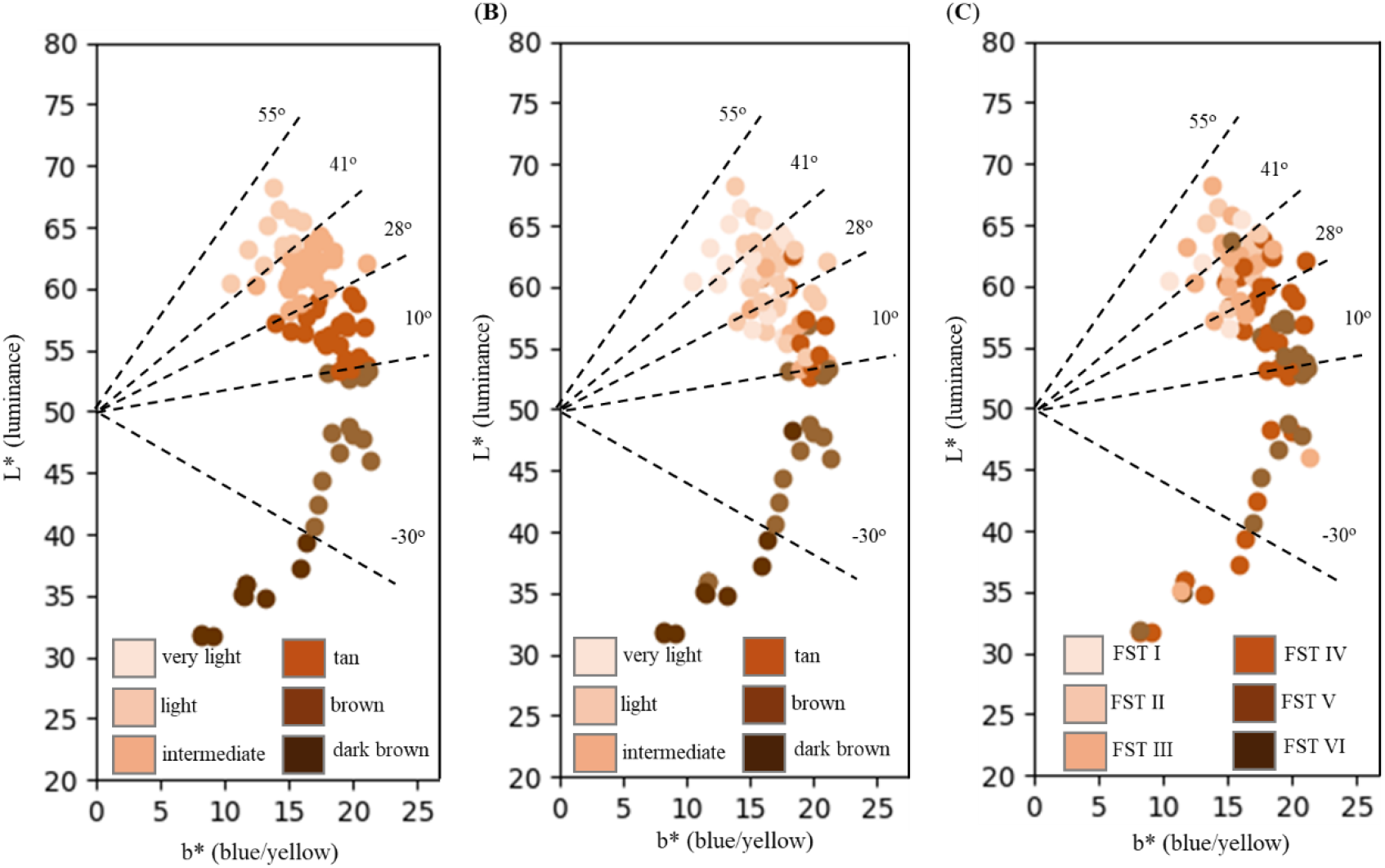
Spread of volunteer skin ITA group. Each dot represents an individual volunteer and placement on the graph represents their objectively determined ITA. ITA bin boundaries are shown as dotted lines. Volunteers are color coded according to their a) objectively measured ITA b) measured PST associated ITA and c) FST.

Figure 2a and b show the spread of the objectively measured volunteer skin spectra, color coded for calculated ITA and for chosen measured PST associated ITA, respectively. Color coding for calculated ITA follows the expected pattern, with the lightest skin having greatest reflectance and the darkest skin the least across the spectrum. Although not a perfect match, PST follows a similar pattern to objectively measured spectra for the darker skin color ITA bins. However, volunteers in the ‘tan’, ‘intermediate’ and ‘light’ ITA groups tend to match their skin to PST swatches that fall into lighter ITA groups. Figure 2c shows measured volunteer spectra color coded for FST and shows that, in general FST is unrelated to spectral color.

**Fig. 2.**
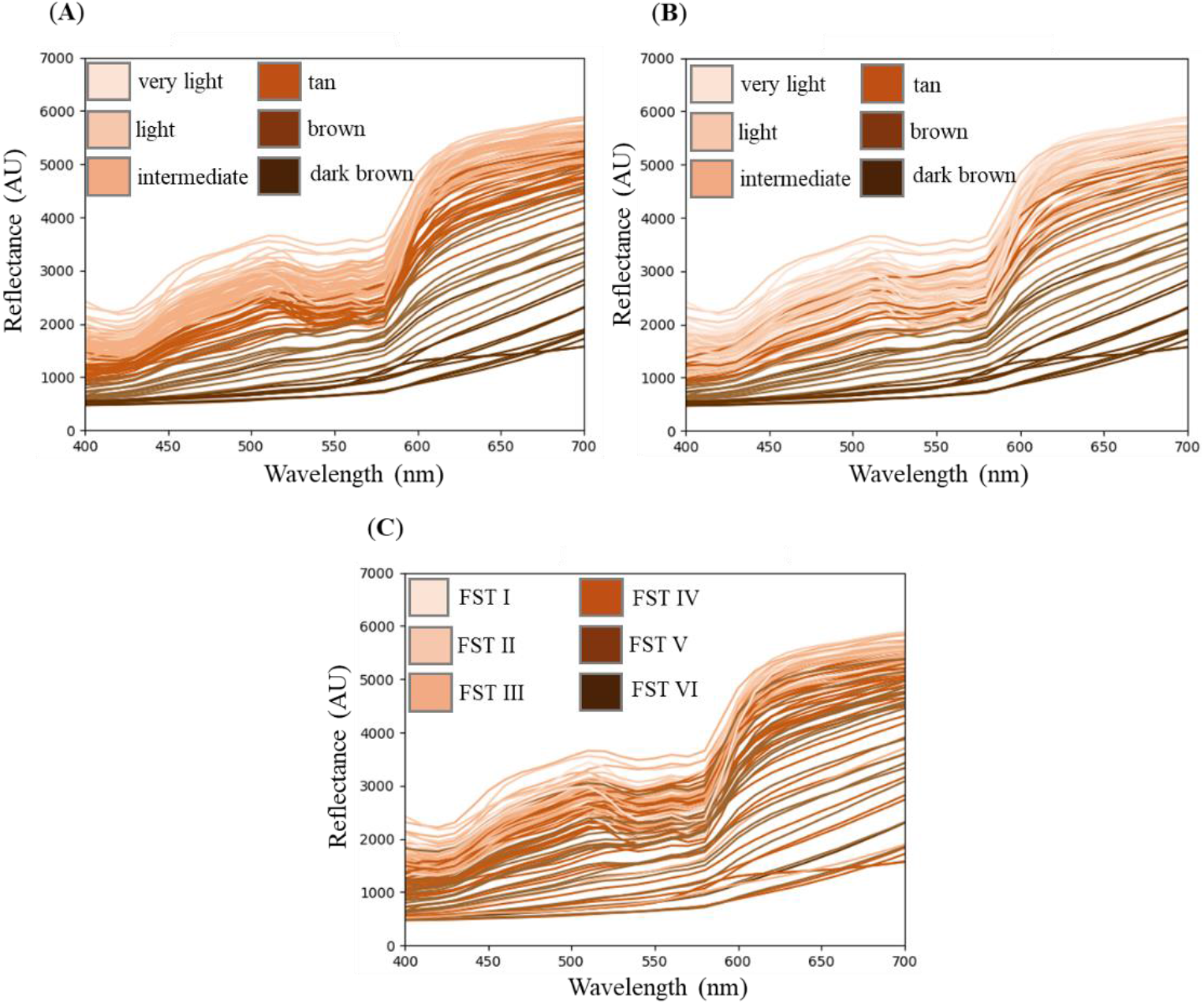
Calculated volunteer skin spectra color coded for a) objectively measured ITA, b) measured PST associated ITA and c) FST.

### 2.3: Correlation between subjective and objective measures

The correlation between the different methods for assessing skin color was calculated using Spearman’s rank correlation or Pearson correlation. Spearman’s rank correlation was used where correlations with FST were calculated. Because measured volunteer and measured PST associated ITA values are both continuous and numerical Pearson correlation could be calculated. For both a correlation coefficient of +/-1 indicates data with a perfect linear correlation, with the sign denoting a positive or negative correlation, and a coefficient of zero implies no correlation. For clinical research, a correlation coefficient of >0.7 is considered useful, but is dependent upon the agreed acceptability[47]. Objectively measured ITA and the subjective FST are negatively correlated with Spearman’s rank correlation ρ = 0.65 (p<0.01). Similarly, PST and FST were negatively correlated with ρ = 0.63 (p<0.01). However, the interquartile range of volunteer data is broader in the darker FST groups (FST IV-V) when skin color is determined using PST compared to objectively measured ITA (Figure 3). Using the PST guide volunteers tend to assess their skin color as being lighter than it is when objectively measured, with most volunteers in FST I, II and III being assigned a PST within the ‘very light’ or ‘light’ ITA group as shown in Figures 1a, 2b and 3b. When skin color is assessed using PST swatches it is determined to be significantly lighter than objectively measured ITA within each FST category, for example volunteers in FST I had an average objectively measured ITA of 38.83° (i.e. ‘intermediate’) but an average ITA associated with PST of 58° (i.e. ‘very light’).

**Fig. 3.**
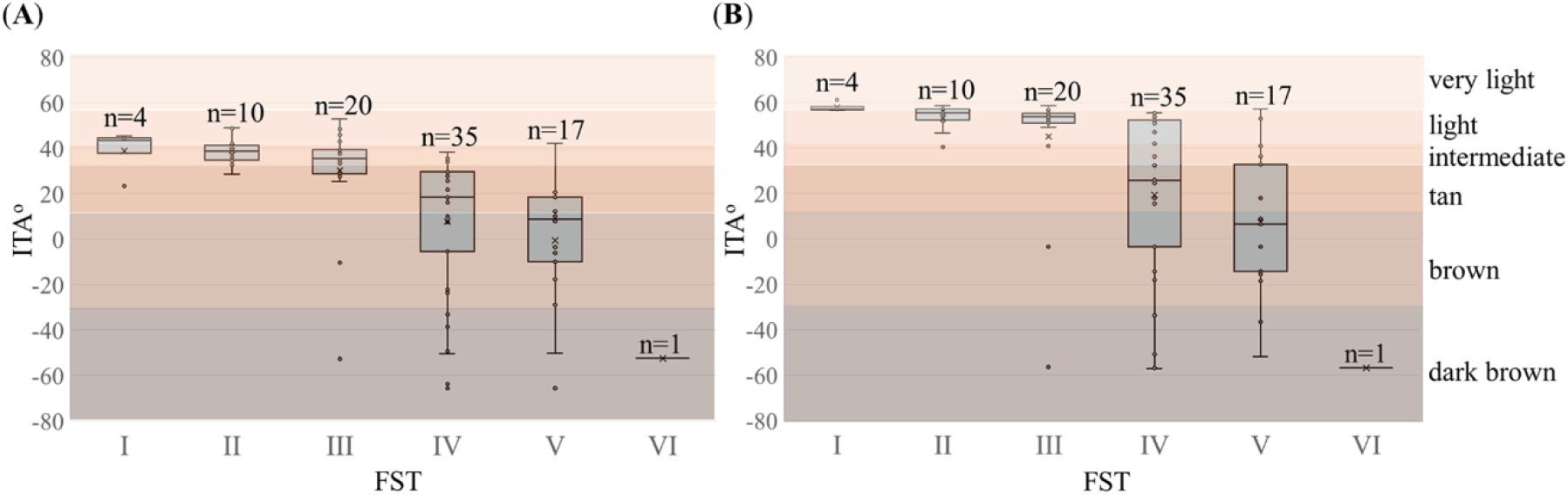
Correlation between objective ITA and FST (a) and measured PST associated ITA and FST (b).

Figure 4 shows the correlation between the objectively measured ITA and the measured PST associated ITA. For a perfect correlation between the objectively measured ITA and the PST associated ITA the ITA color values would match (dotted green line in Figure 4). Although not perfect, correlation is strong with a Pearson coefficient of ρ = 0.90 (p<0.05), therefore the PST guide is a potentially useful subjective replacement for the FST. However, volunteers tend to view their skin as being lighter using PST than when it is objectively measured (p(T ≤ t) = 0.018). Again, the disorder associated with FST, with volunteers color coded for FST, is shown in Figure 4.

**Fig. 4.**
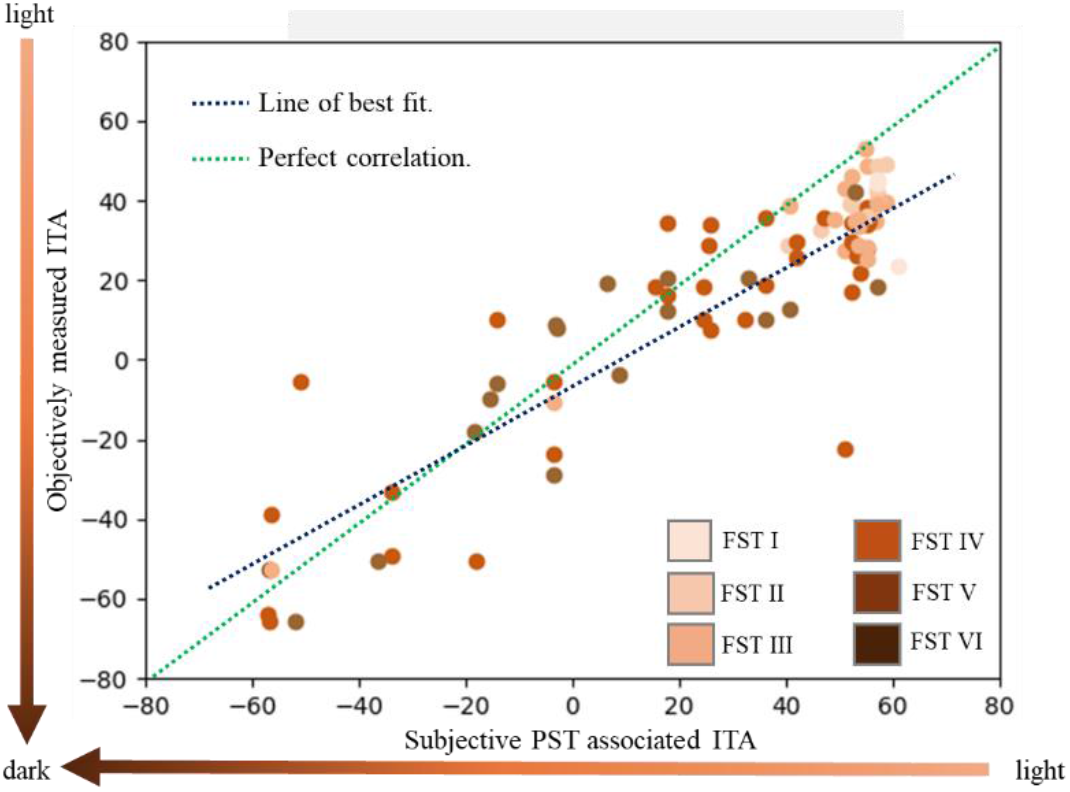
Correlation between objectively measured ITA and subjective PST associated ITA. Volunteers are color coded according to their FST.

### 2.4: Understanding the Monk Skin Tone Scale

As described in section 1.3, the MST has been suggested as a useful alternative to the FST in clinical and research settings[36]. It has recently been recommended for ensuring pulse oximeters are accurate across all skin tones by the FDA, as well as being suggested for use in dermatology settings and research [13, 15, 48]. But how applicable is this scale to healthcare and research in reality? The published CEILAB values for the 10 MST skin color levels were converted to ITA using the formula described in section 1.4[48]. Each color was plotted as shown in Figure 5 for comparison with volunteer data used in both this study and the published data collected by Del Bino and Bernerd from (n = 3,500 volunteers) and Lipnick et al (n = 789 volunteers [24, 25]. Although the total data is dominated by that collected by Del Bino and Bernerd, it provides a good indication of data spread. Combining these data sets is reasonable for this application, particularly since there is good correlation between objective devices and studies[22, 49]. When calculated, MST levels 1-4 all fall into the ‘very light’ ITA category as shown in Figure 5a. Of all skin color measurements used in this study, only 1 volunteer from a total of 4376 fell into this category. MST level 5 is in the ‘light’ skin color ITA group, but it is outside the ITA value of any of the volunteer datasets (Figure 5b). ‘Intermediate’ skin is not represented by MST, and MST level 6 is on the very darkest limit of the ‘tan’ ITA group with an ITA value of 10.81°. The ‘brown’ ITA group is represented by MST level 7. MST levels 8-10 represent the ‘dark brown’ ITA skin color category however, like MST level 5, MST levels 9 and 10 do not represent measured volunteer skin colors.

**Fig. 5.**
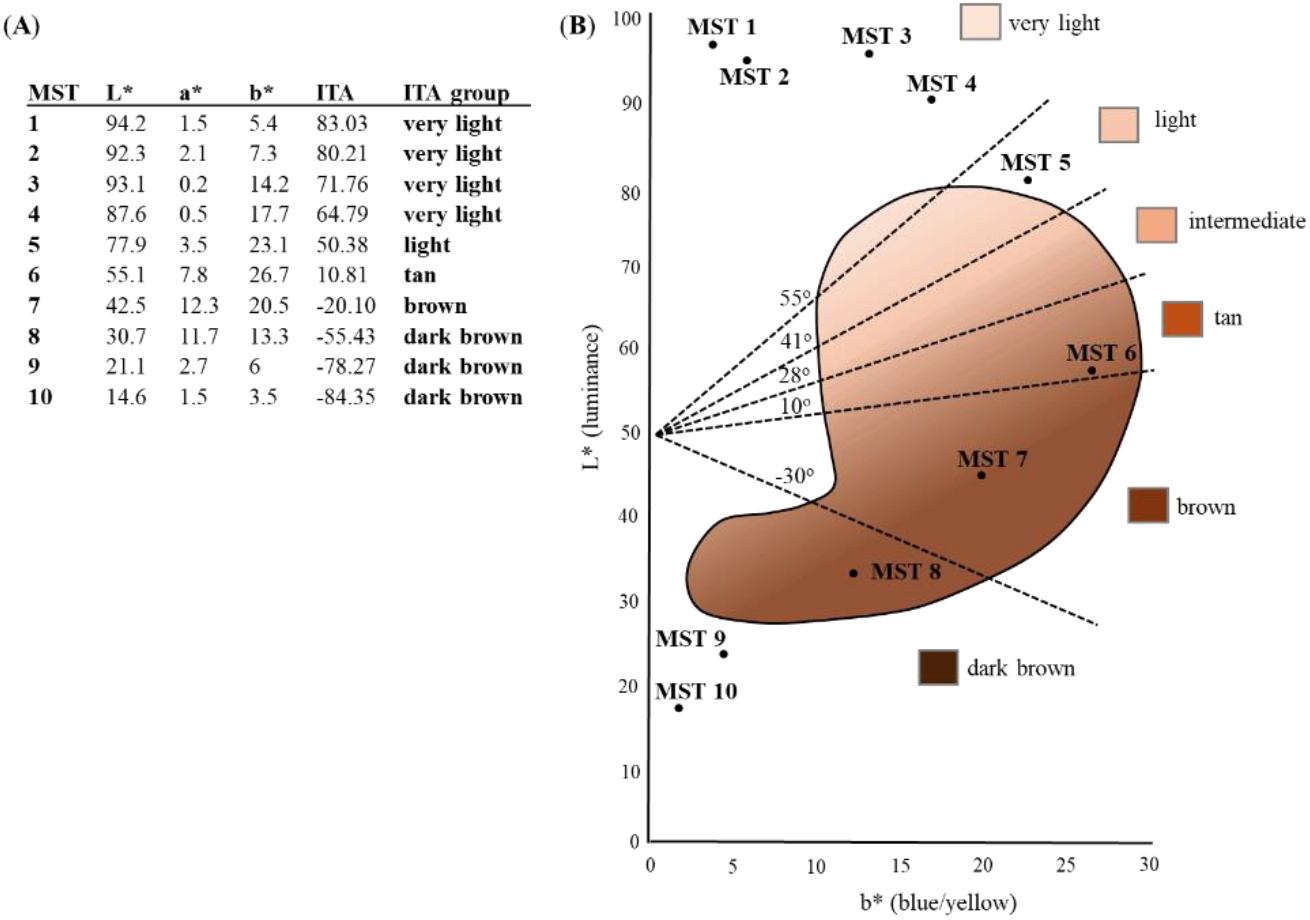
Monk Skin Tone scale. a) MST CEILAB values, ITA calculation and ITA skin color group[50]; b) graphical depiction of MST levels – enclosed area represents the region filled by volunteer skin color measurements including the volunteer cohort discussed in this manuscript (n=87) and from data published by Del Bino and Bernerd (n=3500) and Lipnick et al (n=789) [24, 25].

The MST was created for computer vision and increasing representation in this field. The MST color levels are likely based upon digital images of skin. In a study to create an algorithm to objectively and automatically classify digital skin images, it was shown to be highly accurate, but accuracy was increased using AI generated skin image[15]. However, this method was shown to be equally accurate across all skin tones and accuracy was therefore independent of color.

Whilst useful for computer vision purposes, ambient light during image creation and color scaling of digital images seems to render images too blue/yellow and light/dark. Therefore, as shown in Figure 5, the MST does not represent a realistic spread of skin colors for dermatological purposes. In addition, it has been shown that using printed MST classification for clinical studies may lead to wrongly classifying volunteers. Printers themselves are an issue, with vast variability in color reproduction causing the L* and b* values of the individual MST levels to be significantly skewed from the expected values[37].

## DISCUSSION AND PROPOSALS

Consistent with others, we have shown that FST is not a reliable method for determining skin color for either healthcare or research purposes being highly subjective and being light skin biased with 79% of the volunteers in this study being placed in FST I-IV compared with 72% placed in ITA groups ‘light’ - ‘tan’ (the 4 categories originally created for Caucasian skin for each method)[4, 51–54]. 20% of volunteers in FST I-IV had their skin color objectively measured as being ITA group ‘brown’ or ‘dark-brown’ The FST questionnaire is difficult to interpret with limited and contradictory options, particularly for those with dark skin and up to 60% of individuals are unable to select an appropriate FST questionnaire response for all questions posed, particularly with regard to the ‘reaction to sun exposure’ section of the questionnaire[54]. Volunteers involved in this study found ‘to what degree do you turn tan’ and ‘how sensitive is your face to the sun’ especially difficult to rate, specifically those with ITA < 10° (26.4%). Despite being intended for selecting suitable UV treatment doses in light skinned patients with psoriasis, the FST is still commonly used in practice for unrelated skin color determination including for understanding skin cancer risk[55]. Volunteers in this study belonging to FST groups I-III on average had a very similar ITA values (30° – 39°) meaning that when objectively assessed on average volunteers within FST I-III fall into the ‘intermediate’ individual typology angle (ITA) color category. Volunteers from FST III, IV and V had a large ITA range, suggesting that it is difficult to accurately categorize volunteers into distinct FST color groups (table 2). This was corroborated by volunteer skin color spectra (Figure 2c) which showed that FST is unrelated to spectral color. FST has a negative correlation with ITA (ρ = 0.63; section 2.3) however, the PST was shown to have a very strong correlation with ITA (ρ = 0.9). Therefore, PST would be a useful replacement for FST in dermatology and research being a relatively reliable determinant of skin color and cost-effective.

Lipnick et al., have recently compared a number of methods for determining skin color, including printed versions of the Von Luschan scale and FST and professionally printed MST for direct skin color comparison[25]. Each scale was divided into three groups defined as ‘light’, ‘medium’ and ‘dark’ and these were used to assess the skin color of participants in either ICU with fluorescent lighting or in a research lab under full spectrum lighting. They showed that different methods produce different results when applied to the same cohort, but that the MST ‘dark’ group (consisting of the darkest three shades i.e. shades 8-10) identified volunteers with objectively dark skin better than the other subjective scales tested [25]. Similarly, and in agreement with multiple other authors, we have shown that the FST has limited utility. As discussed in section 2.4 we also show that the MST is not representative of the skin color, and when used as 10 individual colors, rather than the 3 groups described by Lipnick *et al*, does not cover the finer skin color detail required for research and medical device development. However, the subjective PST may provide a useful intermediary between subjective and objective methods due to its much closer correlation with objectively determined ITA than FST. Additionally, the PST chart costs ∼100x less than purchasing a spectrophotometer and is relatively quick and easy to use, not needing to calculate the ITA from the collected spectrophotometer CIELAB data.

As discussed in section 1.4, four ITA skin color groups were initially devised to assess light skin, with two further groups being later added to represent dark skin. The distribution of these groups is not equal and, therefore, makes it difficult to ensure that scientific and medical research is representative. Lipnick et al., raised the question as to how the ITA groups should look to provide equitable representation of the global population[25]. They proposed 2 additional groups to better represent dark skin (taking the total number of ITA color groups to 8) but chose not to split the color groups into equal geometric ITA ranges.

Taking the collective skin color data for a total of 4376 volunteers from this study and those published by Del Bino and Bernerd and Lipnick et al., we propose using 10 skin color ITA groups each separated by 15° (Figure 6)[24, 25]. Due to the significant correlation between PST and ITA, we suggest, for relative ease of use, a set of 9 color swatches (Nottingham Skin Categories – NSC) that represent the border between each of 10 color groups (listed in Figure 6). Roughly half of our volunteers commented on being overwhelmed by the vast number of colors in the 110 swatch Pantone SkinTone Guide (which has recently been updated to consist of 138 shades) and the visual similarity of multiple swatches to their skin color. Therefore, a reduced set of swatches would not only enable easier skin color classification but further reduce costs. Group 1 would represent the lightest skin with an ITA of >55°); group 10 would represent the darkest skin with ITA of < 65°). All groups in between would be separated by 15° and darker than the swatch at the upper boundary, but lighter than the swatch at the lower boundary would be represented by that group. It is difficult to know the proportion of the global population that would be represented by each of these 10 groups. However, based on data collected as part of this study and published by Del Bino and Bernerd and Lipnick et al., a rough estimate of the proportions represented by each group would be: group 1= 0.5%; groups 2 and 6 = 10%; groups 3 and 4 = 19%; group 5 = 15%; groups 7-9 = 7%; and group 10 = 3.5%. If correctly printed or purchased from an approved supplier, although subjective, these could be used cheaply to accurately represent ITA skin color ranges in the clinic and for research purposes (suggested NSC color swatches shown in Figure 6). Compared with the cost of a spectrophotometer of approximately £10,000 GBP, and the requirement for interpretation, these 9 color swatches would be cheap, portable, and simple to use. Used in conjunction with a D65 lamp for skin color determination, these swatches would be suitable for a primary care setting.

**Fig. 6.**
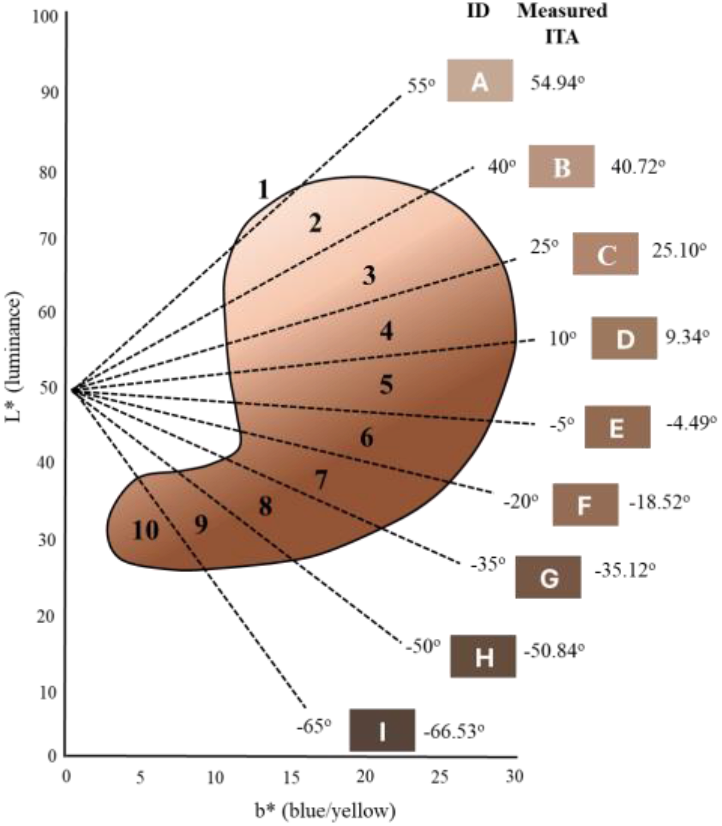
Suggested skin color groupings based on available data, collating the measurements recorded here with Lipnick et al. and Del Bino et al., corresponding to global volunteer skin color[24, 25]. Group 1 would consist of skin with an ITA lighter than 55°(swatch A) and group 10 of skin with an ITA darker than -65°(swatch I). Groups 2-9 are at approximately 15° intervals and bounded by swatches A-I.

To date, most published data regarding skin color has been determined inaccurately and inconsistently typically using FST. Henceforth it is important that this is addressed and that volunteer cohorts for clinical trials of, for example, optical medical devices, are representative. Although objectively determining skin color would be the ideal approach, this is sometimes impractical and expensive. Therefore, we have created a scale with known ITA values, Nottingham Skin Categories, that would be easy to use, without overwhelming the user with a color chart with multiple similar colors. As well as being representative, and categorizing skin color into equally spaced groups, it has potential to provide a relatively accurate understanding of volunteer ITA and increase comparability between studies. However, further work is needed to fully understand the global spread of skin color to ensure inclusivity. The volunteer cohort described here and those collected by other authors have limitations; our cohort and that of Lipnick et al., are from predominantly light pigmented regions and data collated by Del Bino and Bernerd was limited in the locations of the population and gender recruited (e.g. all volunteers were female and African volunteers were recruited from USA and France). Finally, melanin content correlation within our suggested categories would be useful and could be carried out using the methods described by Del Bino and Bernerd, with skin samples from each of the suggested ITA categories.

## MATERIALS AND METHODS

This study was carried out on a cohort of 87 volunteers. Ethics approval was received from the University of Nottingham Ethics Committee to recruit volunteers from within the University. Initially the only limitations placed on demographics was that volunteers were more than 18 years old with no infectious skin disease. However, to ensure to increase numbers of dark-skinned volunteers, Ethics approval was subsequently received to specifically recruit volunteers that considered themselves to have dark skin. Volunteer skin color was objectively determined using the Konica Minolta CM700d spectrophotometer and subjective skin color was determined using the Fitzpatrick Skin Type scale (FST) and the Pantone SkinTone Guide (PST). Results from each were compared with each other and the published L*a*b* values for the MST[48]. A 3mm diameter target mask with plate was used with the spectrophotometer and D65 illumination at 10 degrees to the observer. The specular component was excluded from the measurements, and the selected lens aperture position was small area view (SAV; measurement area) to correctly work with the target mask. The spectrum measured is a ratio of light reflected by the sample at given wavelengths compared to light reflected at the same wavelengths from a calibration sample (certified sample provided by manufacturer). CIELAB color space coordinates were used to calculate the ITA and corresponding ITA skin color group using the formula in section 1.4.

Pantone skin color was determined by holding the guide discussed in section 1.2. against the back of the hand with D65 lamp illumination. The 4-character color code was recorded and photos taken for color matching.

Finally, a Fitzpatrick Skin Type questionnaire was given to each volunteer, as shown in Figure S1, the answers to which were used to determine volunteer FSP. The questions used were those used for skin cancer risk assessment[56]. The scores used to determine FST are shown in Table S1.

## Acknowledgments

This work has been funded by the Engineering and Physical Sciences Research Council, Transformative Healthcare Technologies 2050 (Grant No. EP/T020997/1), InLightenUs.

PhD studentship (SK) funded by EPSRC (Grant No. EP/T517902/1) Studentship award (SK) from SurePulse Medical Ltd

