## Supplementary Material for "Are you represented? Subjective vs objective skin color determination for healthcare and research purposes"

***Fig S1.*** *Fitzpatrick Skin Type questionnaire: self-reported responses are used to determine sensitivity to UV and the total score of the answers to the questionnaire are used to categorize skin color into 6 groups ranging from Type I (fair) to Type VI (dark).*

| Genetic Disposition | | | | | |
|---|---|---|---|---|---|
| **Score** | **0** | **1** | **2** | **3** | **4** |
| **What is the colour of your eyes?** | Light blue, Light grey, Light green | Blue, grey or green | Hazel or light brown | Dark Brown | Brownish Black |
| **What is the natural colour of your hair?** | Red or light blonde | Blonde | Dark blonde or light brown | Dark brown | Black |
| **What is the colour of your skin (non-exposed areas)?** | Ivory white | Fair or Pale | Fair to beige with golden undertone | Olive or light brown | Dark brown or black |
| **Do you have freckles on unexposed areas?** | Many | Several | A few | Very few | None |
| **Total Score for Genetic Disposition** | | | | | |

| Reaction to sun exposure | | | | | |
|---|---|---|---|---|---|
| **Score** | **0** | **1** | **2** | **3** | **4** |
| **What happens when you stay in the sun too long?** | Always burn: Painful redness, blistering and peeling | Often burn: Blistering followed by peeling | Sometimes burn followed by peeling | Rarely burn | Never burn |
| **To what degree do you turn tan?** | Hardly or not at all – I always burn | Light tan | Moderate tan | Tan easily | Turn dark brown quickly |
| **Does your skin tan quickly (within a few hours) after sun exposure?** | Never | Seldom | Sometimes | Often | Always |
| **How sensitive is your face to the sun?** | Very sensitive | Sensitive | Normal | Rarely sensitive | Never had a problem |
| **Total Score for Genetic Disposition** | | | | | |

***Table S1.*** *Fitzpatrick skin type scale score and descriptors.*

The FST groups are described as follows and determined as an answer score from the questionnaire: FST I is very pale skin that burns easily and never tans; FST II is fair skin that burns easily but tans slightly; FST III is medium-colored white skin that burns less easily and forms a gradual tan; FST IV is olive-colored skin that burns occasionally but tans well; FST V is brown skin that rarely burns and tans deeply; and FST VI is dark skin that never burns. FST V and VI are not nuanced enough to categorize brown and dark skin and do not represent the range of darker skin tones[27].

| Score | Description | Fitzpatrick Skin Type |
|---|---|---|
| 0-6 | Highly sensitive, always burns, never tans e.g. very fair skin. | I |
| 7-12 | Very sensitive to sun, burns easily, tans minimally e.g. fair skin. | II |
| 13-18 | Sensitive to sun, sometimes burns, slowly tans to light brown e.g. medium skin. | III |
| 19-24 | Minimal sensitivity to sun, burns minimally, always tans to moderate brown e.g. olive or light brown skin. | IV |
| 25-30 | Insensitive to sun, rarely burns, tans well e.g. brown skin. | V |
| 31+ | Insensitive to sun, never burns, deeply pigmented skin e.g. deeply pigmented dark skin. | VI |

***Fig S2.*** *Pantone SkinTone Guide; 110 skin color swatches each with a central hole for color matching and represented by a 4 digit code describing hue, red/yellow undertone and darkness of skin[31].*

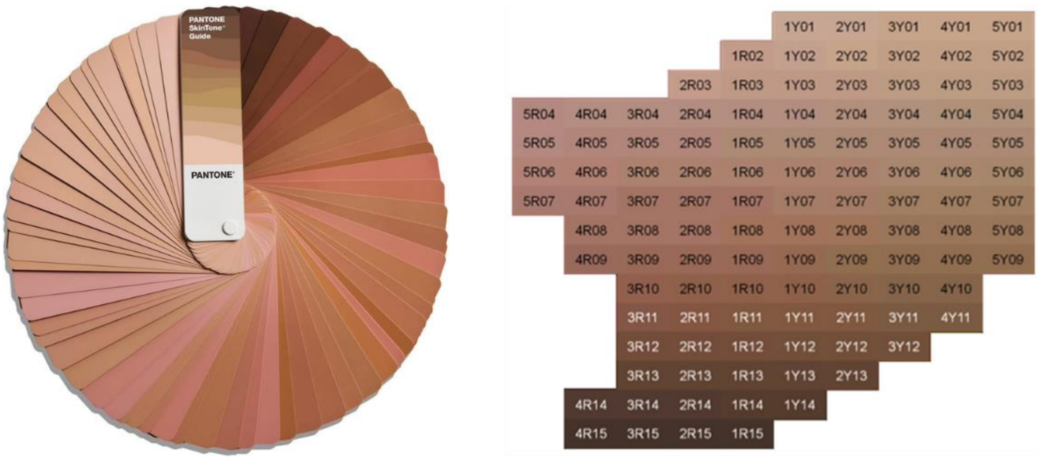


***Fig S3.*** *Monk Skin Tone Scale: the full range of skin tones represented by 10 color levels and intended for use in computer vision applications and optical device testing[20].*

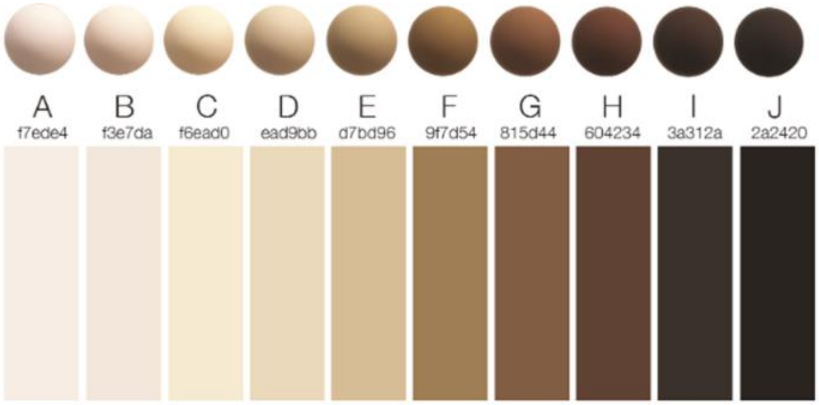


***Fig S4.*** *Skin color classification using Individual Typology Angle (ITA): a. CEILAB color space[43]; b. graphical depiction of skin color as ITA; c. objective ITA skin color bands[48].*

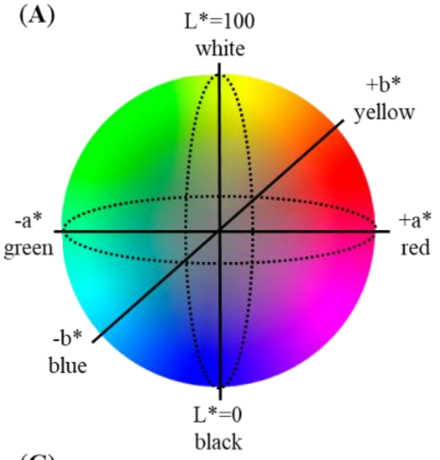


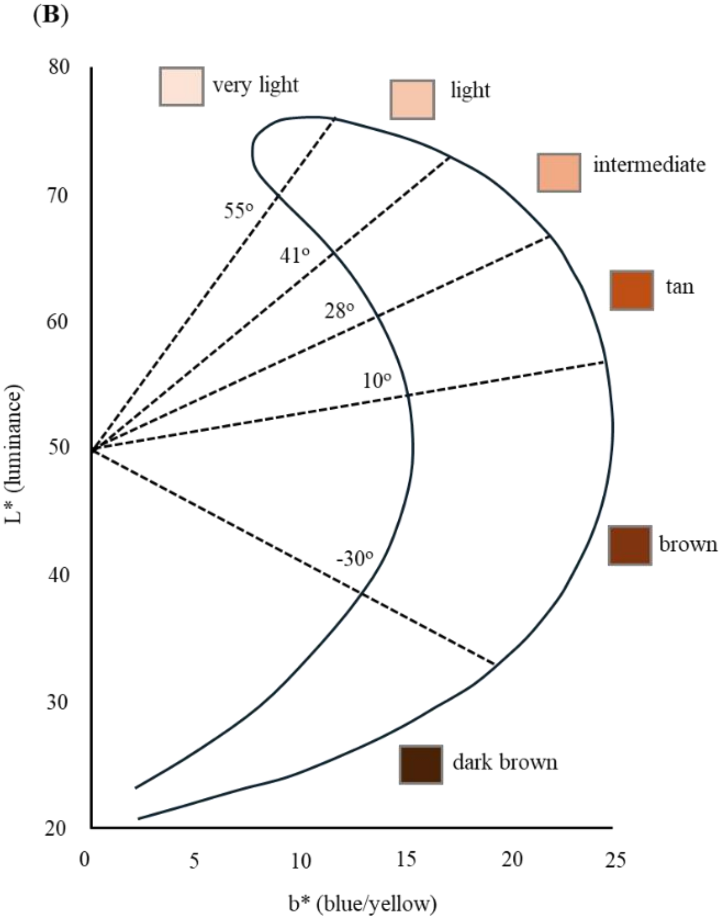


(C)

| **Individual Typology Angle (ITA)** | **Skin color classification** |
|---|---|
| > 55° | Very Light |
| 41° ≤ 55° | Light |
| 28° ≤ 41° | Intermediate |
| 10° ≤ 28° | Tan |
| -30° ≤ 10° | Brown |
| ≤ -30° | Dark Brown |
